# The transcriptional signature associated with human motile cilia

**DOI:** 10.1101/817072

**Authors:** Anirudh Patir, Amy M. Fraser, Mark W. Barnett, Lynn McTeir, Joe Rainger, Megan G. Davey, Tom C. Freeman

## Abstract

Cilia are complex microtubule-based organelles implicated in the aetiology of numerous diseases. Accordingly, many cilia-associated proteins have been described, while those distinguishing cilia subtypes are poorly defined. Here, we characterise the gene signature associated with human motile cilia that captures both known and unknown components of this class of cilia. To define the signature, we performed network deconvolution of transcriptomics data derived from tissues possessing motile ciliated cell populations. For each tissue, genes coexpressed with the motile cilia-associated transcriptional factor, *FOXJ1*, were identified. The consensus across tissues provided a transcriptional signature of 248 genes. For validation, we examined the literature, databases, single cell RNA-Seq data, and the localisation of mRNA and proteins in motile ciliated cells. To validate some of the many poorly characterised genes, we performed new localisation experiments on *ARMC3*, *EFCAB6*, *FAM183A*, *MYCBPAP, RIBC2* and *VWA3A*. In summary, we report a highly validated set of motile cilia-associated genes that helps shape our understanding of these complex cellular organelles.

**Summary:** This work defines a conserved transcriptional signature associated with human motile cilia, including many genes with little or no previous association with these structures. These genes were compared with existing resources and a number of poorly characterised genes validated.

**Graphical abstract:** 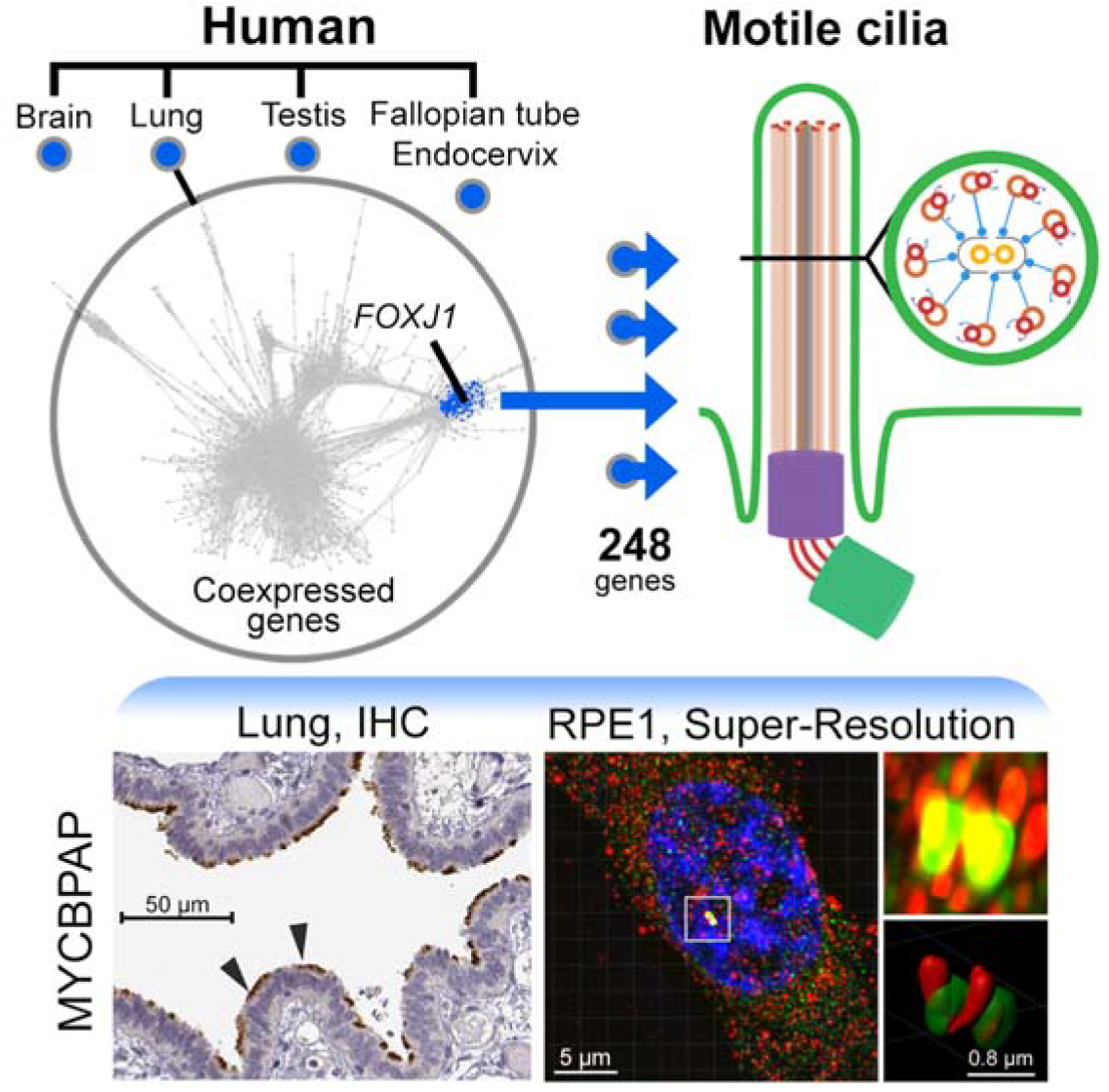

## Introduction

Cilia and flagella are related organelles that facilitate an array of cellular functions. In eukaryotes, the core structural components of cilia includes: the *axoneme*, a microtubular protrusion from the cell surface composed of an array of microtubules; a *centrosomal core*, comprised of a mother (basal body) and daughter centriole (Satir and Christensen, 2007, Mizuno et al., 2012) anchored to the base of the axoneme, and the centriole-associated distal and sub-distal appendages (Uzbekov and Alieva, 2018). Cilia can be subdivided into non-motile primary cilia, in which nine microtubules constitute the axoneme (9+0) and motile cilia, characterised by an additional central pair of microtubules (9+2) (Reiter and Leroux, 2017, Mitchison and Valente, 2017, Hoyer-Fender, 2013). Primary cilia are found on most cell types, where their principal role is as a sensor of the cell’s microenvironment (Anvarian et al., 2019). In contrast, motile cilia are restricted to specific cell populations. Flagellum function as a single large ‘propeller’ and in eukaryotes are found exclusively on spermatocytes where they drive cell motility. Other motile cilia are found in large numbers on the apical surface of certain types of epithelial cells, where their co-ordinated beating displaces the luminal contents over the epithelial surface, e.g. the clearance of mucus in the respiratory tract. Whilst there are a set of core proteins common to all cilia, there are also structural and regulatory elements unique to motile cilia which underpin their distinct functional activity (Heydeck et al., 2018, Choksi et al., 2014b).

Motile cilia play a vital role in human development and homeostasis, and there is a growing list of ciliopathies (cilia-related diseases) associated with mutations of the protein components of these organelles. These include defects in left-right patterning during embryogenesis (Bisgrove and Yost, 2006), infertility (Lyons et al., 2006), asthma (Tilley et al., 2015) and hydrocephalus (Narita and Takeda, 2015). Perhaps the most notable and well-characterised ciliopathy is primary ciliary dyskinesia, an autosomal recessive disorder which has an estimated prevalence of 1 in 10,000 (Praveen et al., 2015, Mitchison and Valente, 2017). Causative mutations leading to primary ciliary dyskinesia include those in genes encoding the motile ciliary components of radial spokes (*RSPH1*, *RSPH9* and *RSPH4A*) (Castleman et al., 2009, Kott et al., 2013), dynein arms, specifically the outer dynein arm (*DNAI1*, *DNAI2* and *DNAH11*) (Guichard et al., 2001, Loges et al., 2008, Bartoloni et al., 2002), and proteins involved in their assembly (*CCDC103, LRRC6* and *ZMYND10*) (Kott et al., 2012, Zariwala et al., 2013, Panizzi et al., 2012). Patients carrying mutations in these genes are often treated for respiratory symptoms, including respiratory infection, due to their inefficient clearance of mucus from the lungs (Mirra et al., 2017, Leigh et al., 2009).

There have already been considerable efforts made to characterise the molecular components of cilia. FOXJ1 and the RFX family of genes have been identified as the key transcription factors which regulate motile ciliogenesis, which in turn, depending on species, have shown to be regulated by the Wnt, Hedgehog and Notch signalling pathways (Choksi et al., 2014b). Moreover, in conjunction with other transcriptional regulators, such as HNF1B and SOX5, further ciliary diversity is introduced for mechanosensory renal cilia and bronchiolar cilia, respectively (Gresh et al., 2004, Kiselak et al., 2010). Proteomic profiling studies have sought to define the components of motile cilia by dysregulating such transcriptional regulators and analysing the proteome of isolated cilia preparations using mass spectrometry (Choksi et al., 2014a, Blackburn et al., 2017, Ostrowski et al., 2002, El Zein et al., 2009, Campbell et al., 2016). Each of these studies has produced a list of cilia-associated proteins and accordingly a number of databases have been established. The most relevant to the study of human cilia include: *CentrosomeDB*, a set of human (and *Drosophila*) genes encoding proteins that are localized in the centrosome, either as centrosome constituents or as centrosome visitors (Nogales-Cadenas et al., 2009); *CilDB* a database dedicated to proteins involved in centrioles, centrosomes, basal bodies, cilia and flagella in eukaryotes (Arnaiz et al., 2009); *SysCilia* a curated list of cilia genes many of which are associated with disease (van Dam et al., 2013); and *CiliaCarta* which employs a naive Bayesian classifier to predict cilia candidate genes across a diverse set of data sets (van Dam et al., 2019). These resources list between 303 and 3376 genes and have greatly broadened our understanding of the complexity of cilia while attempting to define the role of these genes in the context of development, ciliogenesis and ciliopathies.

Here we have sought to provide a consensus human motile cilia gene signature conserved across known motile cilia containing tissues and compare it with the relevant databases. We have used a network deconvolution approach to define gene coexpression clusters containing the transcriptional regulator *FOXJ1* (Yu et al., 2008) using transcriptomics data from the Genotype-Tissue Expression (GTEx) (Lonsdale et al., 2013) project for human tissues known to possess motile ciliated cells. In support of these analyses, we have also examined various lines of evidence in order to validate the set of genes identified. These include a comparison with cilia and centrosomal databases mentioned above, studies of their expression profile across motile and primary cilia containing cells and tissues, and a number of new expression studies examining several poorly characterised genes identified by this work, namely *ARMC3*, *EFCAB6*, *FAM183A*, *MYCBPAP*, *RIBC2* and *VWA3A*. Overall the study proposes a set of motile cilia associated genes that are tightly coexpressed across tissues, including certain but not all cilia-associated centrosomal genes previously identified. The signature genes have been summarised graphically based on their function and/or known localization.

## Results

### Derivation of the human motile cilia signature

The GTEx RNA-Seq (v7) dataset is the largest transcriptomics data resource for non-pathological human tissues currently available and was used here to derive a human motile cilia gene signature. Data derived from tissues likely to contain cell populations possessing motile cilia, i.e. ependymal cells in brain regions likely to adjoin the cerebrospinal fluid-filled ventricular space (n = 863), bronchial epithelia of the lung (n = 427), spermatocytes in testis (n = 259) and tubal epithelial cells in fallopian tube/endocervix (n = 12) were downloaded; in total this represented tissue RNA-Seq data from 1561 samples derived from 566 donors (Figure 1). To identify genes associated with motile cilia, gene correlation networks (GCNs) were generated for each tissue and subjected to cluster analysis. In each case, a gene cluster containing *FOXJ1* and other known cilia proteins was identified (Table S1), ranging in size from 597 to 6126 genes. Such variation in the size of the motile cilia cluster across tissues is likely explained by a varying number of samples and the different tissue biology, e.g. the expression landscape of the testis is dominated by transcriptional signal associated with spermatogenesis, making the flagellum-specific gene module difficult to separate from all sperm-associated gene clusters (Zheng et al., 2019). To take into account the differences in cluster size, we have considered only the 248 gene signature present in all tissue-derived gene lists. However, we acknowledge that the extended list, i.e. 479 genes found in three of the four motile cilia tissue clusters contains many additional validated cilia genes and therefore is likely to contain many other novel motile cilia-associated genes (Table S1).

**Figure 1:**
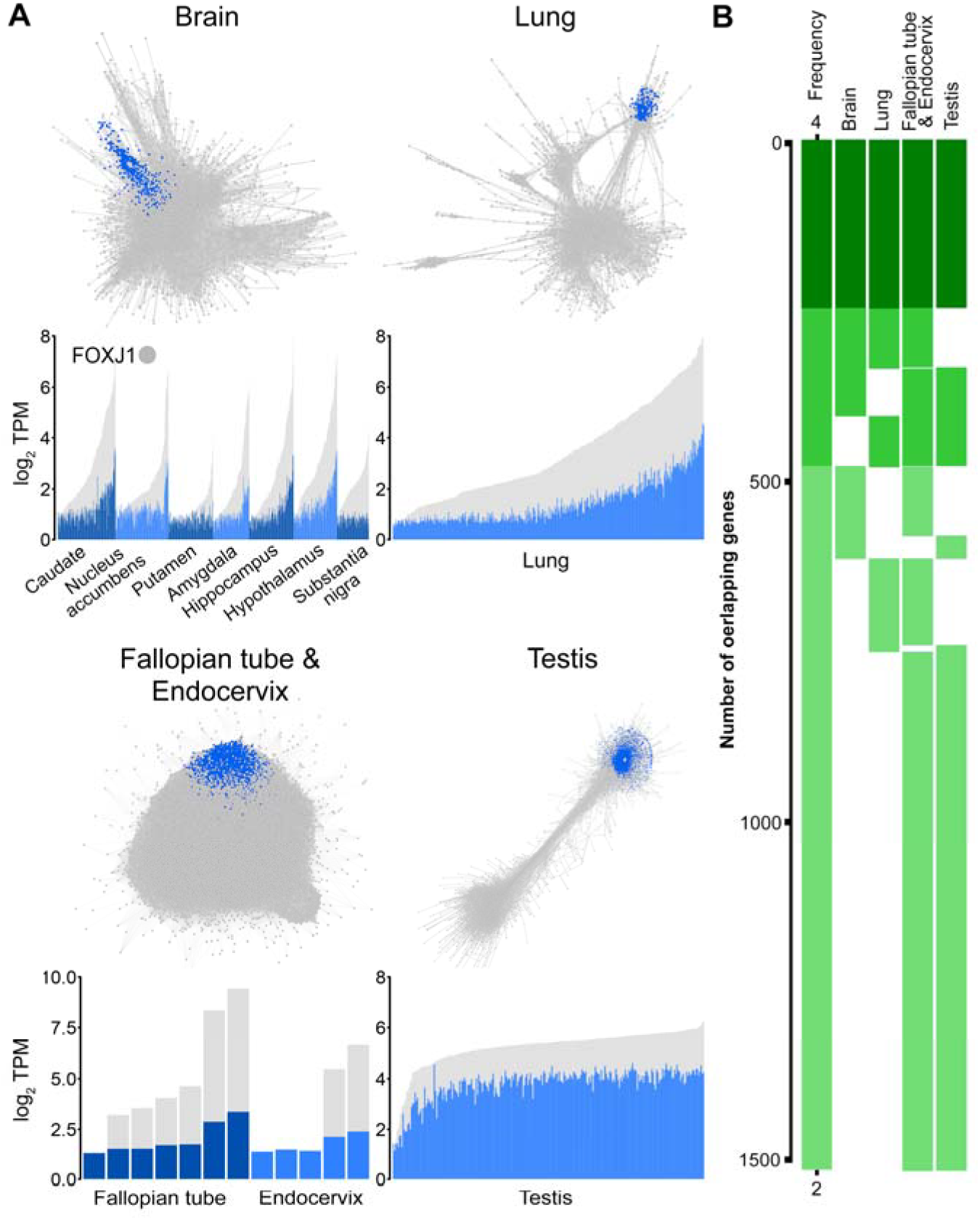
Derivation of the human motile cilia transcriptomic signature. **A)** Gene coexpression networks for four tissue types from the GTEx resource, highlighting genes (blue) co-clustering with *FOXJ1*. **B)** Average expression profile of signature genes (grey) in tissue samples ordered (left to right) based on their expression of *FOXJ1* (blue). **C)** The overlap between the tissue-specific gene clusters sorted from those occurring in all clusters to those shared between two (dark to light green).

### Comparison with database genes and their expression profiles across cells/tissue

Enrichment analysis of the 248 gene signature was conducted for GO terms, pathways, gene families, transcription factor binding sites, and human phenotypes (Tables S2). Enriched gene families included components of the ‘dynein regulatory complex’ (*q* value = 2.2×10^−15^), ‘axonemal, dyneins’, (*q* value = 3.2×10^−13^) and ‘tektins’ (*q* value = 1.4×10^−7^), with the corresponding enrichment of biological processes such as ‘cilium movement’ (*q* value = 3.4×10^−52^), ‘cilium organization’ (*q* value = 5.1×10^−40^) and ‘cilium-dependent cell motility’ (*q* value = 2.7×10^−26^). Additionally, binding sites for *RFX1* and *MIF1* were also found to be enriched for these genes (*q* value < 10^−7^). Human disease phenotypes associated with disorders of motile cilia included ‘abnormal respiratory motile cilium morphology’ (*q* value = 1.2×10^−24^), ‘situs inversus totalis’ (*q* value = 3.4×10^−21^), ‘bronchiectasis’ (*q* value = 1.5×10^−13^), and ‘male infertility’ (*q* value = 3.7×10^−13^). To further investigate the ciliary/centrosomal association of the signature genes we first conducted a literature search on all signature genes. Of the genes identified, the literature supported 133 (54%) as having direct experimental evidence supporting their spatial localization or functional association with cilia (Table S1). A further 87 (35%) genes were found associated with cilia through coexpression analysis but without any direct evidence of their localization within ciliary structures. For 28 genes (11%) no prior association with cilia could be identified.

We then sought to examine the signature’s overlap with public databases of cilia/centrosome proteins. Including the signature reported here, a total of 4333 genes have been implicated previously with cilia and/or centrosomes. These include the CentrosomeDB (Nogales-Cadenas et al., 2009), CilDB (Arnaiz et al., 2009), CiliaCarta (van Dam et al., 2019) and SysCilia (gold standard) (van Dam et al., 2013) (Figure 2A and Table S3). There were only four genes which were common to all the databases and the derived signature (*DNAAF1*, *FOXJ1*, *KIF24*, and *MAK*). In support of our literature search, the majority of the signature genes (196 genes) overlapped with genes listed in CilDB, including well known motile cilia genes such as members of the dynein regulatory complex (*DRC1*, *TCTE1* and *IQCD*), axonemal dynein (*DNAH2*, *DNALI1* and *DNAI1*) and tektin gene family (*TEKT1*, *TEKT2*, and *TEKT4*), whilst also including genes with poor evidence supporting an association with cilia, e.g. *MYCBPAP*, *ARMC3* and *EFCAB6*. Relative to the databases, 52 genes were found to be unique to the current study and included genes not associated with human motile cilia previously, e.g. *FAM183A* and *VWA3A*. By contrast, 84 genes recorded by all database resources were absent from the derived signature. Upon inspection, these largely represented genes associated with the cell cycle (Giotti et al., 2018) and ciliary assembly and maintenance, e.g. members of the centrin family (*CETN1*, *CETN2* and *CETN3*), BBSome complex members (BBS1, BBS4, BBS5 and BBS7) and IFT genes (*IFT20*, *IFT74* and *IFT81*) (Lechtreck, 2015).

**Figure 2:**
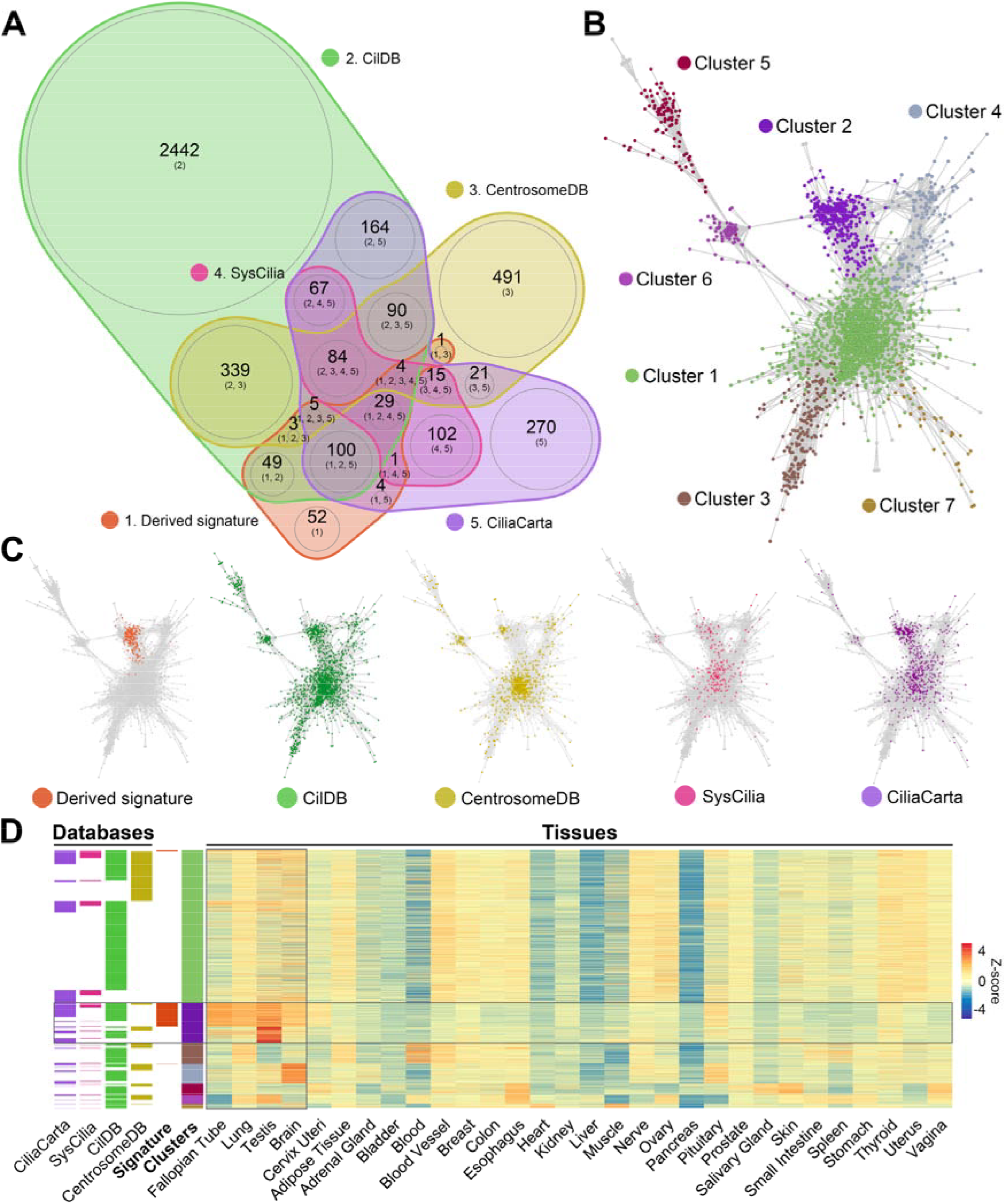
Comparison of signature and database genes and their expression across tissues. **A)** Venn diagram of overlapping genes between the derived signature and databases using nVenn (Pérez-Silva et al., 2018). Here the overlap between gene lists can be observed through the overlap of their respective segments and based on the numbers shown in brackets, pointing to the numbered gene lists. **B)** Gene coexpression network for these genes across the GTEx tissues, highlighting genes from the clusters. **C)** Expression Z-score of the signature and database genes across GTEx tissues ordered based on the clustering from **B**).

As a further analysis, we examined the global expression patterns of all signature genes and those recorded in databases, examining their expression across all 51 tissue types in the GTEx resource. GCN analysis was again used to visualise and explore the expression profile of signature and database genes across human tissues (Figure 2B-D). Cluster analysis was used to broadly group genes together based on their underlying expression pattern (Figure 2B). Highlighting the genes from each database, showed them in each case to be distributed across the network. In contrast to the distribution of signature genes were far more localized. This is indicative of their tight coexpression across all tissues (Figure 2C), attributed by their relatively high expression in tissues known to have motile ciliated cells (Figure 2D). Conversely, genes for each of the databases were scattered throughout the graph suggesting that the genes had very different expression profiles, ranging from broad expression across all tissues as represented by cluster 1, to highly expressed in certain tissues such as blood (cluster 3) or brain (cluster 4). Additionally, cluster 3 included many immune genes, e.g. associated with MHC class 1 and 2, TLR receptors and TNF family of genes (Table S3). As a more direct comparator, the expression of signature genes was examined in single cell RNA-Seq data derived from the mouse brain and lung (Figure 3). Here motile ciliated ependymal and bronchial cells, respectively, showed a significantly higher average expression of signature genes (*q* value < 0.001) when compared to other cell types, again supporting their specific association with motile cilia possessing cells.

**Figure 3:**
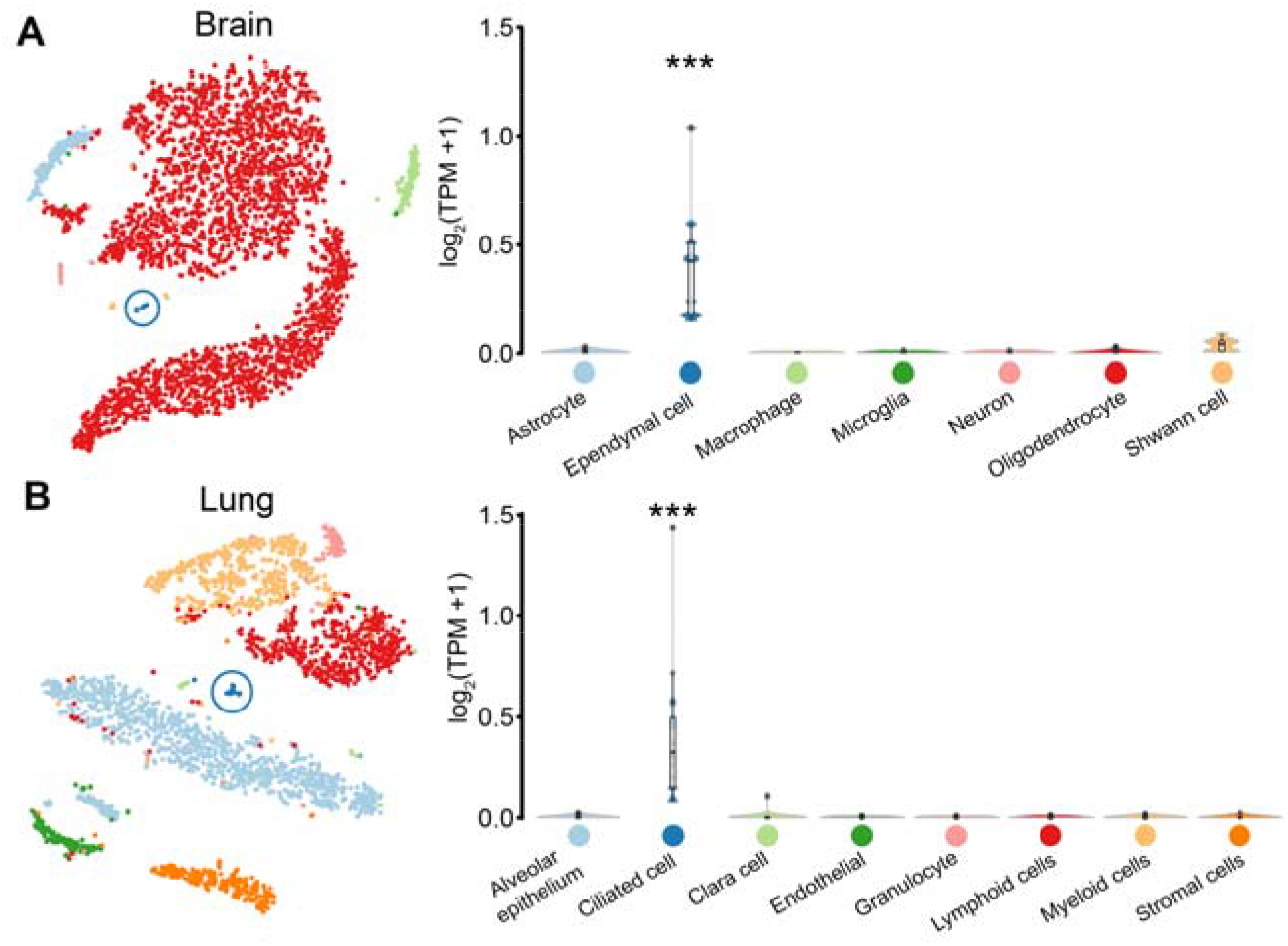
Expression of signature genes in cells from the brain and lung. Average expression of signature genes across cells of the **A)** brain and **B)** lung taken from the mouse cell atlas dataset (Han et al., 2018).

### The localisation of candidate proteins in motile ciliated tissue

In order to provide an additional level of validation for the 248 signature genes, we examined the immunohistochemistry (IHC) data in the Human Protein Atlas (HPA) resource (Uhlen et al., 2010) in tissues containing motile ciliated cells (Figure 4, Table S1). Based on this, the genes were placed into three groups: high confidence genes (n=119, 48%) were those where positive staining for the cilia/centrosome was observed in at least one tissue with no staining of other structures. Medium confidence (n=50, 20%) was assigned to genes where the protein was positively stained for in cilia/centrosome, but the data also showed staining of other structures. Finally, for 79 (32%) genes no data was available or no apparent staining was observed on the sections, and they were designated as being unsupported by this approach. In no cases, did we observe any evidence of the specific staining of non-ciliated cells.

**Figure 4:**
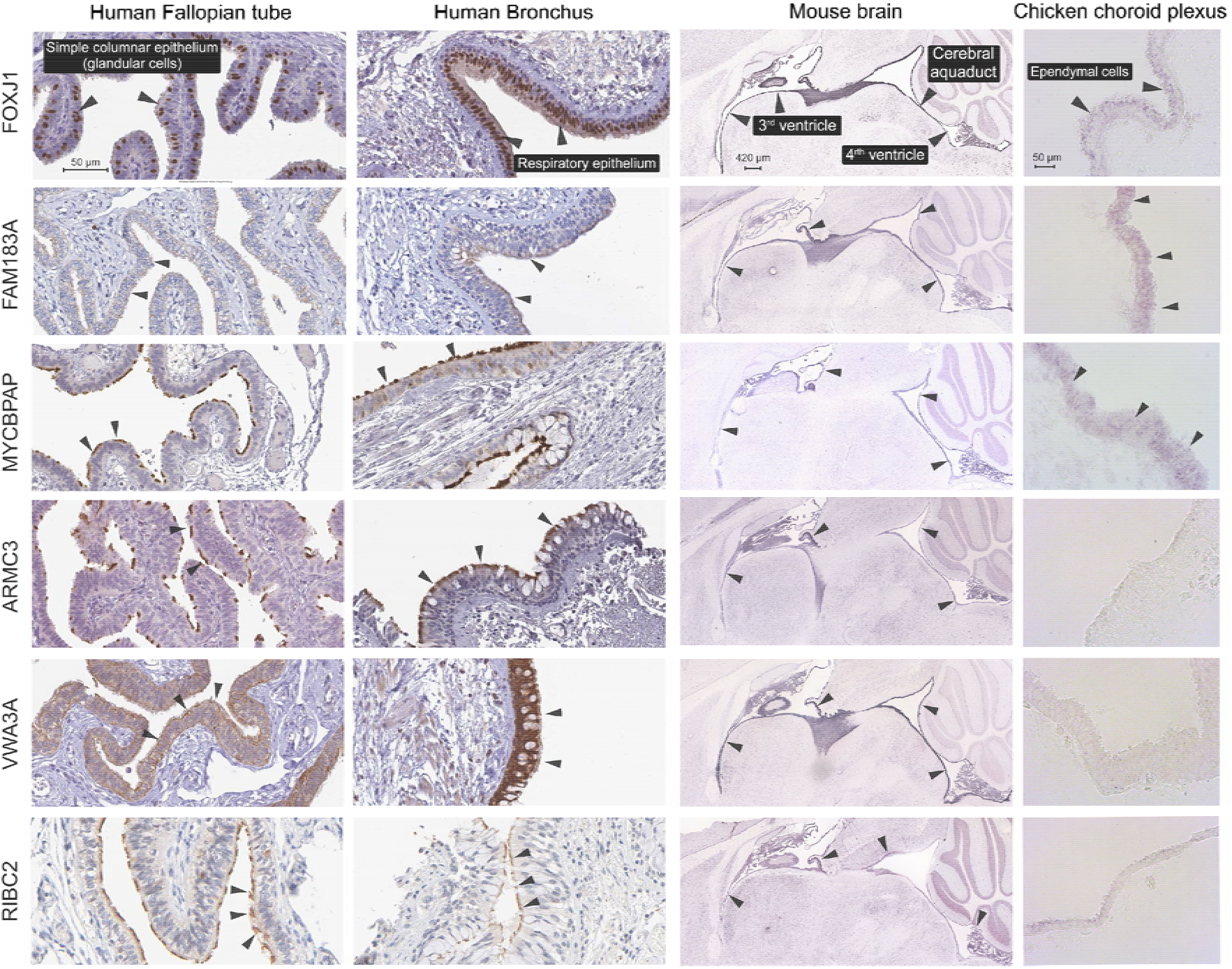
IHC and ISH staining from signatures genes across species and tissues. IHC and ISH staining of tissue sections from the HPA and Allen brain atlas of the mouse brain for encoded proteins and transcribed RNA of signature genes, respectively. The final column, new ISH staining performed in choroid plexus sections from chicken embryos (stage 35).

### Experimental validation of uncharacterised cilia-associated genes

After collating the results of the above analyses, six genes with little evidence in the literature of an association with cilia were selected for further investigation. This included five high confidence genes based on our assessment of the HPA IHC data: *ARMC3, FAM183A, MYCBPAP*, *RIBC2* and *VWA3A*, and *EFCAB6* which had no HPA data associated with it. For these genes, we examined localised gene expression through RNA *in situ* hybridisation (ISH) data from the Allen mouse brain atlas (Sunkin et al., 2013). Furthermore, we performed further ISH analyses on sections of the choroid plexus from chicken embryos (stage 35) looking for staining in ependymal cells (Figure 4 and S1) which have motile cilia (Stephen et al., 2013). In all cases, positive staining for motile ciliated cells was observed in mouse brain, however, ISH staining of ependymal cells lining the choroid plexus in the chickens was only observed in the cases of *EFCAB6, FAM183A and MYCBPAP*. Next, we sought to see if ARMC3, MYCBPAP, RIBC2 and VWA3A were associated with the primary cilia of human RPE1 cells. IHCs were conducted in cells at G_0_ and M phase, using confocal microscopy while those in G_0_ were further examined by super-resolution microscopy (Figure 5). Confocal imaging showed MYCBPAP and RIBC2 localised to the centrosomes in both phases, VWA3A localised to the centrosome only in M phase, ARMC3 showed no colocalization (Figure S2). Super-resolution microscopy further revealed, MYCBPAP and RIBC2 co-localising with both the mother and daughter centriole in G_0_ phase.

**Figure 5:**
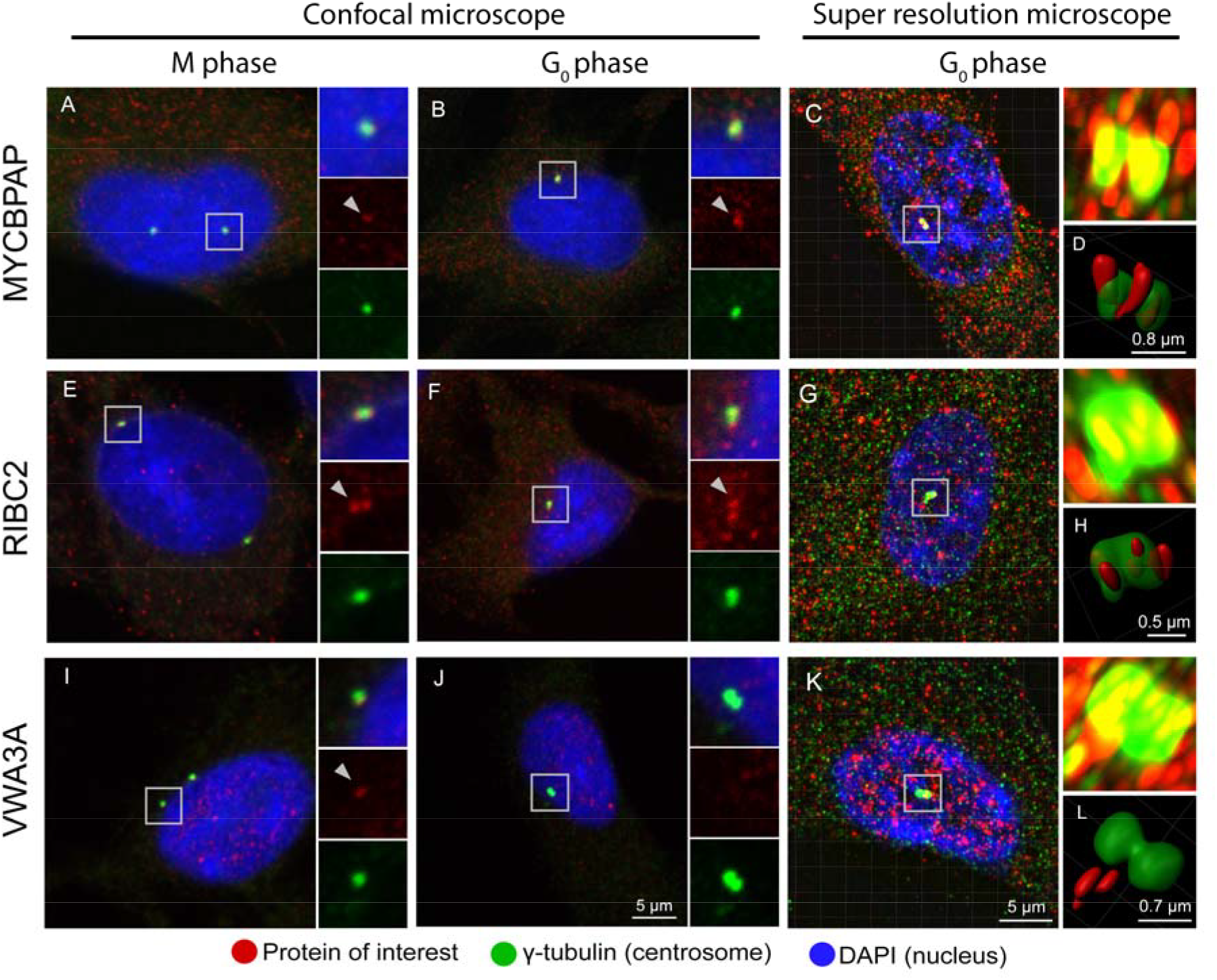
Localization of encoded proteins in RPE1 cells. IHCs for encoded proteins (red) of signature genes in RPE1 cells at M (**A**, **E** & **I**) and G_0_ (**B-D**, **F-H** & **J-L**) phase, with centrosomal marker y-tubulin (green) and nuclear marker DAPI (blue). Confocal imaging shows centrosomal colocalization of MYCBPAP (**A**, **B**), and RIBC2 (**E**, **F**) with y-tubulin in both phases, while the same localization is seen for VWA3A only in M phase (**I**). Super-resolution microscopy at G_0_ confirms MYCBPAP (**C**) and RIBC2 (**G**) localization. With added 3D modelling in Imaris (**D**, **H** & **L**), images show their broad localization towards both mother and daughter centrioles.

As a graphical summary of this work, we sought to categorize all motile cilia signature genes based on their known association with cilia (Figure 6, Table S1). The figure broadly categorized genes into different levels of confidence based on literature mining, including those with a known localization (grey box), cilia-association but no localization (green box) and those with no association at all (yellow box). Genes examined experimentally here, are highlighted in blue.

**Figure 6:**
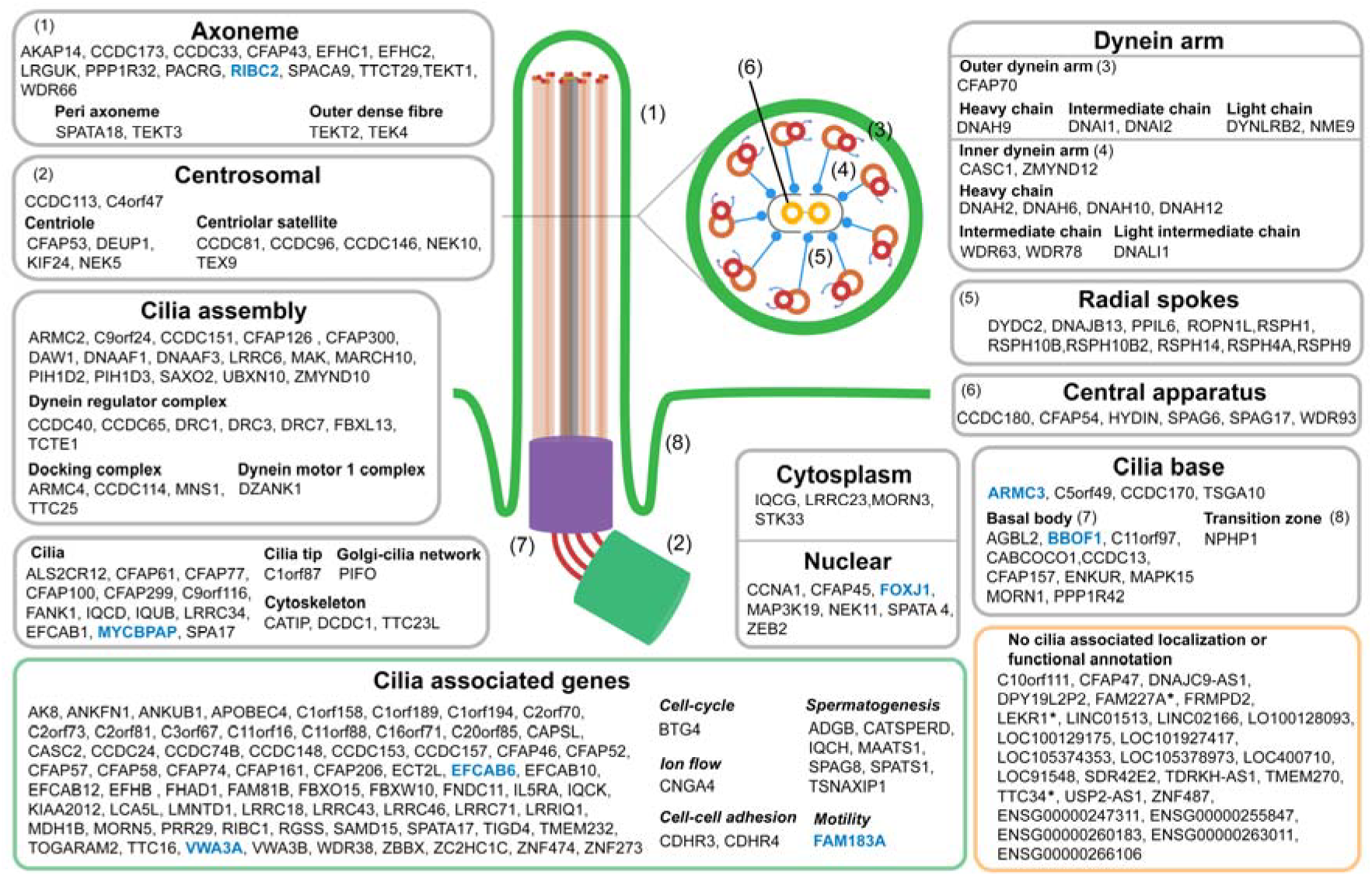
Annotation of derived signature genes. Annotation of signature genes based on previous literature in the context of cilia, giving priority for the spatial localization to their function. Genes having evidence for cilia associated localization are grouped in grey boxes. Genes associated with cilia, however lacking any localization evidence are grouped in the orange box. While those without any association with cilia are grouped in the yellow box, with high or medium confidence genes based on our analyses marked with an asterisk. Additionally, genes examined in this study are highlighted in blue.

## Discussion

Motile cilia are a distinct class of cilia. They are characterized by a 9+2 configuration of central microtubules, radial spokes and dynein arms, along with specialized proteins involved in their assembly. Together, the microtubules and associated molecular motors allow cilia to beat in an ATP-dependent manner (Sanchez et al., 2011). They are vital in cell motility, embryonic patterning, fertilization and the movement of luminal contents over epithelial surfaces. Mutations in the components of these organelles result in a wide range of disorders termed as ciliopathies. Studies in humans and model organisms have identified thousands of proteins as potentially being involved in the cilia biogenesis, maintenance and assembly, and the results of these studies are presented in a number of databases, i.e. CentrosomeDB (Nogales-Cadenas et al., 2009), CilDB (Arnaiz et al., 2009), CiliaCarta (van Dam et al., 2019) and SysCilia (gold standard) (van Dam et al., 2013). One of the challenges in identifying candidate genes or proteins specific to motile cilia is the fact that many components are involved in other cellular processes or structures. For example, centrosomal replication is associated with cell division, during which many components are upregulated (Giotti et al., 2018), and primary cilia, which contain many of the same proteins, are present in most cell types.

Here we have attempted to harness the power of GCN’s and employ the principle of ‘guilt-by-association’ to identify genes specifically associated with motile cilia. This is based on the fact that genes specifically associated with a given cell type or biological process, frequently vary in expression with their relative abundance or activity within a sample and consequently across a large sample set the expression of these genes is tightly correlated. This approach has been used previously to identify genes associated with specific cell populations and processes, from tissue and cell-level transcriptomics data (Mabbott et al., 2013, Patir et al., 2019, Nirmal et al., 2018, Shih et al., 2017). Here we analysed the brain, lung and the female reproductive tract, all of which contain populations of multi-ciliated cells which function to move luminal contents (cerebrospinal fluid, mucus) over the epithelial surface. In addition, we examined the testis, where cilia proteins are associated with the flagellum of sperm, a fundamentally different type of motile cilia but comprised of many of the same molecular components. In the current study, we first identified genes from each of the selected tissues which co-clustered with *FOXJ1*, a key transcriptional regulator of the motile ciliogenic program (Yu et al., 2008, Vij et al., 2012). In the case of the brain and lung, a clear transcriptional module associated with multiciliated epithelial cells was defined due to the marked variation in the abundance of these cell populations across the samples. For the female reproductive tract and testis, however, such modules were harder to define accurately, as there were either only a few samples available and with flagellum-related genes being strongly associated with genes involved in spermatogenesis (Zheng et al., 2019), respectively. To circumvent these limitations and filter out any cell type-specific genes, we compared the gene clusters from each tissue to arrive at a consensus signature of 248 genes. It should be noted, however, that the list of genes associated with three of the tissue clusters (an additional 231 genes) also contained many other known cilia proteins and therefore by inference genes encoding other uncharacterised cilia components (Table S1).

Validation of the gene signature included enrichment analyses, annotation based on a literature review and cilia-associated databases, and exploration of other resources describing the cellular expression of genes and proteins, confirmed the majority to be known components of motile cilia or associated regulatory systems (summarised in Figure 6). The binding site for the transcriptional factor *RFX1*, a member of the RFX gene family (Piasecki et al., 2010), was enriched. This gene has shown to be involved in development, based on a mouse knockout model and regulates the basal body-associated protein *ALMS1*, defects in which cause Alström syndrome ciliopathy (Feng et al., 2009, Purvis et al., 2010) but was not present in the signature. However, *RFX3* was found in three of the tissue-derived motile cilia signatures, making it a likely candidate as a regulator for motile-ciliogenesis, an observation supported by a recent study (Chen et al., 2018). In addition, the transcriptional binding site for MIF was also enriched in signature gene promoters. MIF is known to affect cell motility through the regulation of microtubule formation (Pick et al., 1979, Winner et al., 2008).

The signature genes were also cross-referenced with the four cilia/centrosome gene databases; CentrosomeDB (Nogales-Cadenas et al., 2009), CilDB (Arnaiz et al., 2009), CiliaCarta (van Dam et al., 2019) and SysCilia (gold standard) (van Dam et al., 2013) databases. The majority (79%) of genes in the signature were corroborated by one or more of the databases. Notably, many well-known primary cilia genes involved in ciliary assembly and signalling and listed by the databases were absent from the signature. This included members of the BBSome complex, IFT chain (Wei et al., 2012, Ye et al., 2018) and many associated with the centrosome. As known components of primary cilia, these genes are regulated through the stages of cell cycle and are ubiquitously expressed across cell types, and would therefore be expected to have a different expression profile in the tissues examined relative to the genes coexpressing with *FOXJ1*. To explore the expression of signature and database genes across tissues, GCN analysis was used for all the 51 tissues from the GTEx project. Interestingly, although the signature was derived from separate analyses of individual tissues, in general their coexpression was highly conserved across the 51 tissue types, being highly expressed in motile ciliated tissues relative to others. Interestingly, these also included a number of centrosomal genes indicative of a specialised centrosomal system for motile cilia assembly and function. Genes listed by the various databases coexpressing with those of the signature included known motile cilia components like dyneins (*DNAH3*, *DNAH7*, and *DNAH8*) and members of the α-tubulin gene family (*TUBA3D*, *TUBA3E*, and *TUBA3C*) (Konno et al., 2015, Fischer et al., 2009). In contrast, analysis of databases showed genes within a given database to be distributed across the GCN, and exhibit little evidence of co-expression, suggestive of representing different biology across tissues. Closer inspection showed some to be immune-related genes, e.g. TLR and MHC genes, and their presence in the databases is likely an artefact of the approaches used to define them (Ross et al., 2007). As a direct validation of their specificity of expression, single-cell transcriptomics data derived from the mouse brain and lung showed the signature genes to be highly and specifically expressed in ependymal and ciliated epithelial cells, respectively, of these tissues.

The HPA resource was used to further verify the validity of signature genes based on IHC analysis. Genes for which there was data were ranked as being of either high or medium confidence based on their expression pattern matching that expected for proteins associated with motile cilia. Our analysis of the HPA data showed it to validate the majority of signature genes; 48% were scored as high confidence genes and 20% as medium confidence, based on the criteria outlined in the methods. Nothing could be concluded for the 32% genes for which no data was available or the data was of very poor quality. Remarkably, none of the protein localisation data directly contradicted an association of any gene with motile cilia. We then set out to further investigate six genes, *EFCAB6* having no HPA data and five high confidence genes based on their HPA IHC results but with poor evidence of their association with cilia in humans based on literature: *ARMC3*, *FAM183A*, *MYCBPAP, RIBC2* and *VWA3A*. ISH experiments performed on the chicken choroid plexus, showed *FAM183A, EFCAB6 and MYCBPAP* to be expressed by motile ciliated cells lining this tissue, although this was not apparent for *RIBC2*, *VWA3A* and *ARMC3*. Apart from being a possible false negative, this discrepancy could be indicative of the diversity of ciliary components in eukaryotes (Nevers et al., 2017). A recent study using evolutionary proteomics has predicted *MYCBPAP* to be associated with cilia, and ISH of the *FAM183A* orthologue has shown to positively stain motile ciliated tissue in *Xenopus laevis (Beckers et al., 2018, Sigg et al., 2017)*. Additionally, *EFCAB6* has previously been associated with sperm motility in human through proteomics (Amaral et al., 2014). In support of our observations, ISH data from the Allen brain atlas for the selected genes showed positive staining of ciliated cells lining the ventricles of the mouse brain. Further localization experiments of ARMC3, MYCBPAP, RIBC2 and VWA3A were conducted to examine protein expression during G_0_ and M phase, as ciliogenesis is dictated by the different cell cycle stages. RIBC2 and MYCBPAP were observed to colocalise to centriole in both phases and the sub-cellular localization of VWA3A, which to our knowledge has not been examined previously, showed it to colocalise to the mother and daughter centriole in M phase. No colocalization was observed for ARMC3 within the primary ciliated RPE1 cells, although our examination of tissue levels IHC from the HPA and Allen brain atlas did show motile cilia staining for this protein. In the case of *RIBC2*, which is known to function within the cell cycle (Giotti et al., 2018), the gene also influences motile ciliary beating in *Chlamydomonas*, where it has shown to colocalized to the axoneme (Chung et al., 2014). Such findings of multifunctional genes add to the emerging literature on the alternative regulation of centrosomal and cell cycle genes for motile- and multi-ciliogenesis or associated with their function, as also suggested by their coexpression with *FOXJ1* across tissue from the GTEx (Vladar et al., 2018, Zhao et al., 2013, Balestra and Gönczy, 2014). Indeed, studies have found certain cell cycle genes uniquely regulated through alternative promoter usage depending on the cell type (Giotti et al., 2018). Finally, we have sought to summarise our findings graphically, based on a literature search of the known associations of the signature genes with cilia structures. Clearly, many of the known components of the motile cilia machinery have been identified by this study, and many others have evidence supporting their association but not with specific components of the organelle. The curation of the list clearly highlights the many potentially novel cilia genes/proteins identified by this work.

In summary, we have used coexpression analyses to identify a set of 248 genes highly associated with the presence of motile ciliated cells within human tissue. Significant efforts were then made to validate the genes identified based on further coexpression analyses, extensive searches of the literature, online resources of information on the cellular and tissue expression data for genes and proteins, as well as public databases of cilia related genes across different species. Along with a graphical description of signature genes within cilia, the signature highlights similar genes from cilia and centrosome databases, helping in the categorization of known cilia genes towards their role in motile-, primary-or multi-ciliated cells. In the case of a number of poorly described genes we identified, i.e. *ARMC3*, *EFCAB6, FAM183A, MYCBPAP*, *RIBC2* and *VWA3A* we have been able to provide new evidence supporting their association with motile cilia. Such analyses serve to extend and refine the list of genes/proteins specifically associated with motile cilia, allowing more targeted analyses of their localisation and functional role within these complex and important organelles.

## Material and methods

### Data pre-processing, signature derivation

Pre-normalized RNA-Seq data from the GTEx project (Lonsdale et al., 2013) was downloaded (version 7) and log-transformed. Data for tissues known to possess motile ciliated cells were sub-sampled. These included samples taken from seven regions of the brain (n = 863), lung (n = 427), testis (n = 259), fallopian tube (n = 7) and endocervix (n = 5). Due to the small number of samples of fallopian tube and endocervix, data from these tissues were combined. As such, the relative content of motile cilia containing cells varied considerably across samples, with the expression of genes specifically associated with these structures varying accordingly. Motile cilia-associated genes were identified for each individual tissue by GCN analysis. In order to generate a GCN, a gene-to-gene Pearson correlation matrix was calculated between all genes using the network analysis software, Graphia (Kajeka Ltd., Edinburgh, UK). A threshold of *r* ≥ 0.8 was then applied such that only genes correlated to others above this threshold were connected by an edge. In each case, a structured GCN was generated with modules of coexpressed genes forming highly connected cliques within the network. These were defined as clusters using the Markov clustering algorithm (MCL) (van Dongen and Abreu-Goodger, 2012), using an inflation value MCLi = 2.2 (which defines the granularity of clustering). Putative motile cilia-associated genes were defined as those present in the same cluster as *FOXJ1.* Accordingly, four gene clusters were obtained, one for each tissue type. This approach has been adopted previously to identify co-regulated genes with a related function or associated with a given cell type (Nirmal et al., 2018, Shih et al., 2017, Patir et al., 2019). From the four tissue-derived signatures, those genes common to all four signatures were considered for the final human motile cilia signature. Evidence for an association with cilia was explored through literature mining and enrichment analysis, conducted using ToppGene (Chen et al., 2009).

### Functional annotation of motile cilia signature genes and comparison with databases

Evidence for an association of the 248 motile cilia signature genes with cilia was explored through literature mining and enrichment analysis, conducted using ToppGene (Chen et al., 2009). Signature genes were then compared to genes listed in the databases of cilia and centrosomal components, i.e. CentrosomeDB (Nogales-Cadenas et al., 2009), CilDB (Arnaiz et al., 2009), CiliaCarta (van Dam et al., 2019) and SysCilia (gold standard) (van Dam et al., 2013) were collated based on their Ensembl gene IDs and compared to the signature list derived here (Table S3). The expression profile of this combined list was examined across 51 tissues (excluding samples derived from pooled cells) of the GTEx dataset (n = 11,215, donors = 713). A GCN was then generated using these genes only, again using a correlation threshold of *r* ≥ 0.8 and the resultant graph was clustered using a low inflation value (MCLi = 1.2), so as to provide a coarse grain segmentation of the graph comprising of 17 clusters.

To explore the expression of the signature genes at a cellular level, single cell transcriptomics data from the mouse brain and lung were taken from the Mouse Cell Atlas (Han et al., 2018) and analysed. These tissues were selected as they include populations of motile ciliated cells. The batch corrected expression matrices based on unique molecular identifiers were downloaded from the mouse cell atlas database (https://figshare.com/articles/MCA_DGE_Data/5435866). This included batch 1 of the brain (n = 3285 cells) and lung (n = 2501 cells) cell data. Additionally, for the latter, 11 cells annotated as “dividing cells” were excluded, as it was unclear which cell types these referred to. Corresponding mouse orthologues for signature genes were identified based on their Ensembl gene ID using BioMart (Hubbard et al., 2002). The average expression of signature genes was then tested for significance in the ciliated cell populations (ependymal cells of the brain and ciliated epithelial cells of the lung), versus all other cell types as defined in the mouse cell atlas. The non-parametric Wilcoxon signed-rank test was adopted for these comparisons.

### Immunohistochemistry and RNA in situ hybridisation

The tissue distribution of mRNA and proteins for all signature genes were investigated using publicly available resources. IHC staining of human tissue sections from the bronchus and fallopian tube were examined in the HPA (Uhlen et al., 2010). In both tissues, positive staining of the ciliated epithelial cells lining the tissue was considered as validatory evidence.

The expression of a number of novel genes were further examined in the choroid plexus of chicken embryos (stage 35, day 9) by ISH. All use of animals was undertaken in accordance with the Animals (Scientific Procedures) Act 1986, UK. For selected genes, clones which covered the majority of exons near the centre of the gene were preferentially selected (*ARMC3*: ChEST208k22, *EFCAB6*: ChEST912jB, *FAM183A*: ChEST261m5, *FOXJ1*, *MYCBPAP*: ChEST864g6, and *RIBC2*: ChEST770c15) using the UCSC Genome Browser (Karolchik et al., 2003) and where available obtained (Source BioSciences, UK) (Boardman et al., 2002). Fertilised chicken eggs were incubated for nine days at which point the embryos were sacrificed, the choroid plexus dissected and tissues fixed overnight in 4% paraformaldehyde (PFA) at 4°C. Samples were then rinsed in PBS and equilibrated overnight in 15% sucrose/PBS before embedding in sucrose-gelatin (15%:7.5%) and snap frozen in isopentane at −70°C. Cryostat sections (10 µm) were cut and stored overnight at −20°C. Sections were then rinsed in PBS and fixed for overnight in 4% PFA. After successive rinses with PBS, the tissue was permeabilized by incubation in proteinase-K (20 ng/ml K-03115836001 Roche) for 10 min at room temperature. Sections were treated consecutively with 4% PFA, acetic anhydride solution (0.25% acetic anhydride and 1.3% triethanolamine) with intermittent washing. Finally, 5 nM probe in hybridisation buffer (50% formamide, 5xSSC pH 4.5, 0.05 µg/ml yeast RNA, 0.05 µg/ml heparin, and 1% SDS) was applied to the slides. Following an overnight hybridization with probes at 65°C, sections went through a series of post-hybridization washes and then maleic acid buffer-tween (0.15 M NaCl, 0.1 M maleic acid, 0.18 M NaOH and 0.02% tween). After blocking (20% heat-inactivated FBS/KTBT) for 1 h, sections were incubated overnight with 1:1000 anti-digoxigenin-alkaline phosphate (11093274910 Roche, 1:1000) at 4°C. Following a final series of washing with MABT, sections were incubated with staining solution (3.5 µl/ml nitro blue tetrazolium, N-6876 Sigma; 3.5 µl/ml BCIP, B-8503). After staining for 1-2 h (depending on the probe), the reaction was stopped by several rinses in PBS. ISH staining of brain sections for these genes was also examined in data from the mouse Allen brain atlas (Lein et al., 2006), where the choroid plexus and ventricular system were present (which is lined with motile ciliated ependymal cells).

### Confocal and super-resolution imaging

RPE1 cells were arrested in G_0_ and M phase followed by immunolabeling and imaging. To arrest cells in M phase they were treated with 1:100 KaryoMAX Colcemid solution (15212012; Gibco, UK) in PBS for 3 h. To arrest cells in G_0_ and induce ciliogenesis, cells were serum starved for 24 h. Cells were then fixed with methanol at −20°C for 10 min. After washing in PBS, cells were immunolabeled with polyclonal antibodies against ARMC3 1:100 (HPA037824, Sigma), MYCBPAP 1:100 (HPA023257, Sigma), RIBC2 1:100 (HPA003210, Sigma) and VWA3A 1:100 (HPA041696, Sigma), and a monoclonal antibody against gamma-tubulin 1:1000 (T5326, Sigma) prior to incubation with appropriate secondary antibodies conjugated to Alexa Fluor 488 (A11017, Invitrogen) and Alexa Fluor 594 (A21207, Invitrogen). Finally, samples were stained with DAPI and mounted for confocal imaging on the Zeiss LSM710 Confocal Microscope.

Super-resolution images (SIM) were acquired using structured illumination microscopy. Samples were prepared on high precision cover-glass (Zeiss, Germany). 3D SIM images were acquired on an N-SIM (Nikon Instruments, UK) using a 100x and 1.49 numerical aperture lens with refractive index matched immersion oil (Nikon Instruments). Samples were imaged using a Nikon Plan Apo TIRF objective (numerical aperture 1.49, oil immersion) and an Andor DU-897X-5254 camera using 405, 488 and 561 nm laser lines. Z-step size for Z stacks was set to 0.120 µm as required by the manufacturer’s software. For each focal plane, 15 images (5 phases, 3 angles) were captured with the NIS-Elements software. SIM image processing, reconstruction and analysis were carried out using the N-SIM module of the NIS-Element Advanced Research software. Images were checked for artefacts using the SIMcheck software (http://www.micron.ox.ac.uk/software/SIMCheck.php). Images were reconstructed using NiS Elements software v4.6 (Nikon Instruments, Japan) from a Z-stack comprising of no less than 1 µm of optical sections. In all SIM image reconstructions, the Wiener and Apodization filter parameters were kept constant.

## Supplemental material

**Table S1: Human motile cilia associated signature and evidence, with overlapping genes across tissue derived signatures**.

The derived motile cilia signatures for each of the selected tissues from the GTEx project (Lonsdale et al., 2013) using gene coexpression network analysis. This includes the frequency of overlapping genes across these tissue derived signatures. Additionally, the human motile cilia signature, with supporting evidence and annotation. This includes evidence from the various databases, and the HPA, as well as the annotation as described in Figure 6 and their ranking based on the HPA analysis (Uhlen et al., 2010).

**Table S2: Functional enrichment analyses for signature genes**.

Results from enrichment analyses based on ToppGene (Chen et al., 2009).

**Table S3: Comparison of signature and database genes, and their GCN analysis across all GTEx data**.

Comparison of signature and database gene lists. Moreover, the clustering of these genes in GTEx using gene coexpression networks.

**Figure S1: IHC and ISH staining for EFCAB6.** ISH staining of *EFCAB6* transcribed RNA in tissue sections from the mouse brain (Allen brain atlas) and those performed in choroid plexus sections from chicken embryos.

**Figure S2: Localization of ARMC3 in RPE1 cells.** IHCs of ARMC3 (red) in RPE1 cells at M (**A**) and G_0_ (**B-D**) phase, with centrosomal marker y-tubulin (green) and nuclear marker DAPI (blue). The co-localization experiments were examined using confocal imaging (**A**, **B**) and super-resolution microscopy (**C**) at G_0_ with added 3D modelling in Imaris (**D**) showing broad localization towards the mother and daughter centrioles.

## Acknowledgements

T.C.F., M.G.D., M.W.B., J.R. and L.M. receive funding from an Institute Strategic Programme Grant awarded by the Biotechnology and Biological Sciences Research Council, Institute Strategic Program Grant [BB/P013732/1: ‘BluePrints for Healthy Animals’]. A.M.F. is supported by a BBSRC EastBio Studentship.

## Author contributions

*Anirudh Patir*: Data curation; Formal analysis; Investigation; Validation; Visualization; Writing - original draft; Writing – review & editing

*Amy M. Fraser*: Investigation; Validation; Writing – review & editing

*Mark W. Barnett*: Validation

*Lynn McTeir*: Validation

*Joe Rainger*: Methodology; Validation; Writing – review & editing

*Megan G. Davey*: Conceptualization; Funding acquisition; Resources; Supervision; Writing – review & editing

*Tom C. Freeman*: Conceptualization; Funding acquisition; Resources; Supervision; Writing – review & editing

## Abbreviations

GCN: Gene coexpression network
GTEx: Genotype tissue expression
IHC: Immunohistochemistry
ISH: In situ hybridization

