## Supplementary figures and images for "The transcriptional signature associated with human motile cilia"

### Figure S1

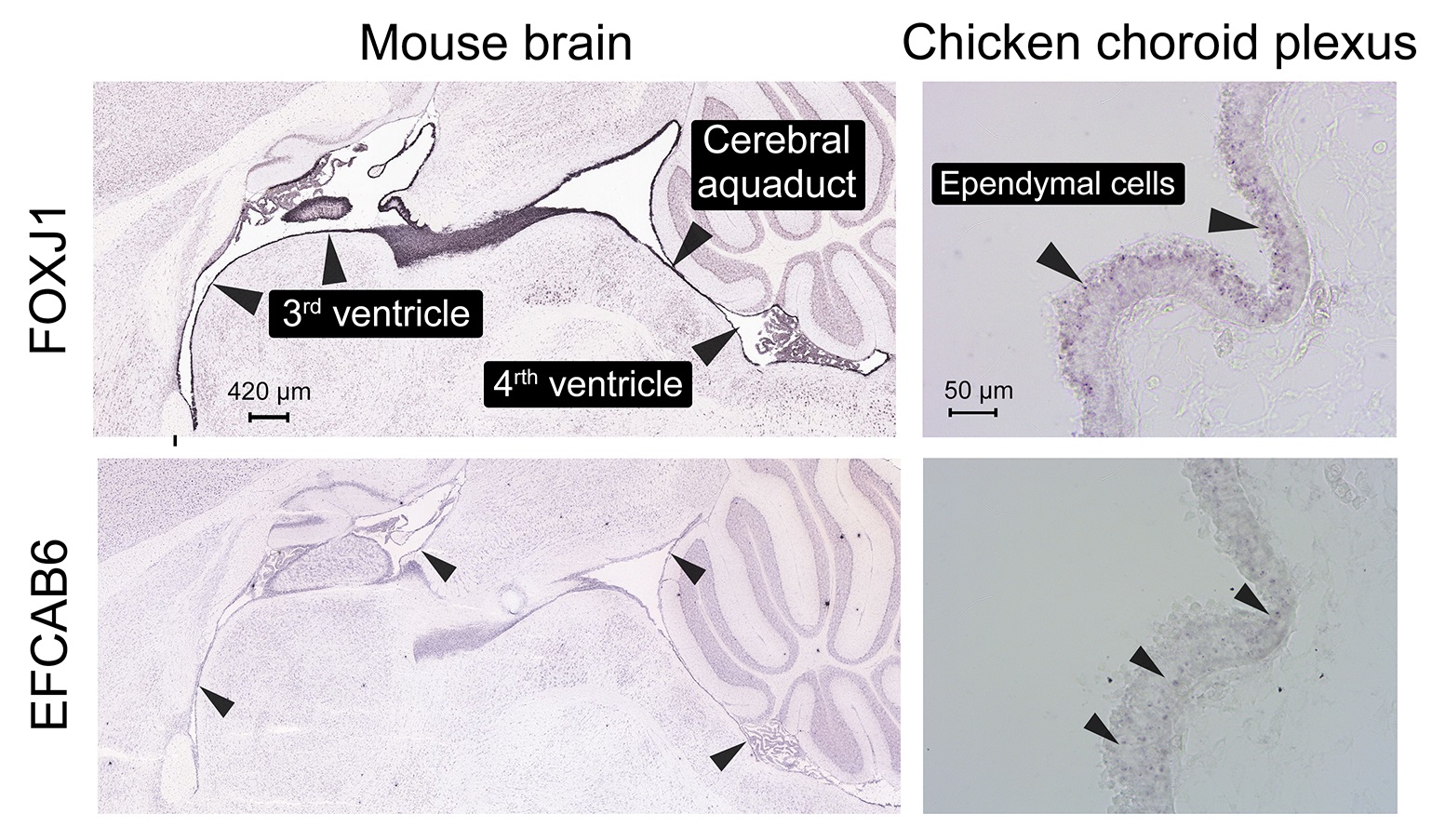

### Figure S2

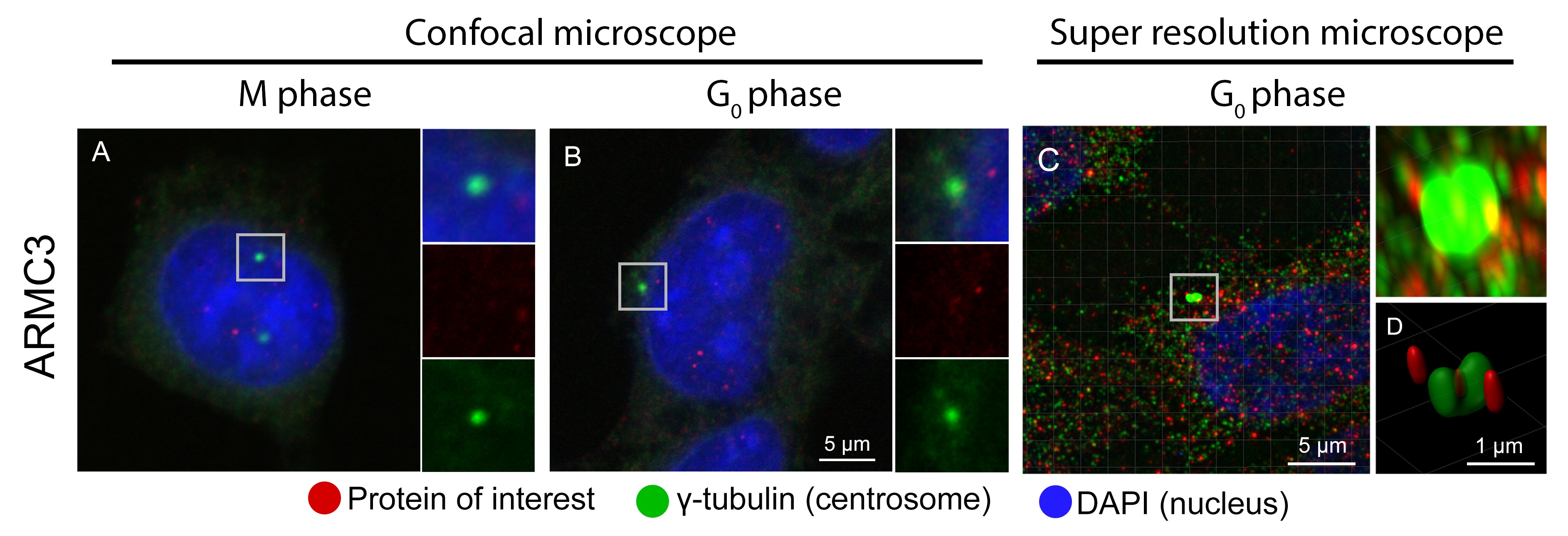
